# Stress affects the epigenetic marks added by natural transposable element insertions in *Drosophila melanogaster*

**DOI:** 10.1101/037598

**Authors:** Lain Guio, Cristina Vieira, Josefa González

## Abstract

Transposable elements are emerging as an important source of cis-acting regulatory sequences and epigenetic marks that could influence gene expression. However, few studies have dissected the role of specific transposable element insertions on epigenetic gene regulation. *Bari-Jheh* is a natural transposon that mediates resistance to oxidative stress by adding cis-regulatory sequences that affect expression of nearby genes. In this work, we integrated publicly available data with chromatin immunoprecipitation experiments to get a more comprehensive picture of *Bari-Jheh* molecular effects. We showed that *Bari-Jheh* was enriched for H3K9me3 in nonstress conditions, and for H3K9me3, H3K4me3 and H3K27me3 in oxidative stress conditions, which is consistent with expression changes in adjacent genes. We further showed that under oxidative stress conditions, H3K4me3 and H3K9me3 spread to the promoter region of *Jheh1* gene. Finally, another insertion of the *Bari1* family was associated with increased H3K27me3 in oxidative stress conditions suggesting that *Bari1* histone marks are copy-specific. We concluded that besides adding cis-regulatory sequences, *Bari-Jheh* influences gene expression by affecting the local chromatin state.

## INTRODUCTION

Gene regulation is a complex process that involves mechanisms at the DNA sequence level and at the epigenetic level. Although genes can acquire novel regulatory mechanisms through different types of mutations, transposable elements (TEs) are emerging as an important source of regulatory variation ^1,2^. TEs can contain cis-regulatory sequences that affect the expression of nearby genes. Some of the recent examples on the global impact of TEs on gene expression levels include: providing enhancer sequences that contribute to the stress-induced gene activation in maize, adding transcription factor binding sites in the mouse and the human genomes, and providing alternative transcription start sites in Drosophila ^3-5^. The epigenetic status of TEs can also affect gene regulation. In *Arabidopsis thaliana*, gene transcription is affected by the methylation status of intragenic TEs ^6^ and correlates with siRNA-targeting of TEs ^7^. In Drosophila, local spreading of repressive heterochromatin marks from TEs has been associated with gene down-regulation ^8,9^. Although all these studies strongly suggest that TEs may play a role in gene regulation through different molecular mechanisms, detailed analyses that link changes in expression with fitness effects are needed to conclude that TEs have a functional impact on gene expression.

There are a few examples in which TE-induced changes in gene expression have been shown to be functionally relevant ^10-12^. One of these cases is *Bari-Jheh,* a *Drosophila melanogaster* full-length transposon providing a cis-regulatory sequence that affects the expression of its nearby genes ^13,14^. *Bari-Jheh* is associated with downregulation of *Juvenile hormone epoxy hydrolase 2* (*Jheh2*) and *Jheh3* in nonstress conditions, and with upregulation of *Jheh1* and *Jheh2* and downregulation of *Jheh3* under oxidative stress conditions ^12^. We have previously shown that *Bari-Jheh* adds Antioxidant Response Elements to the upstream region of *Jheh2* leading to *Jheh2* and *Jheh1* upregulation under oxidative stress conditions ^12,15^. However, how *Bari-Jheh* affects gene expression under nonstress conditions, and how *Bari-Jheh* affects *Jheh3* expression under oxidative stress conditions remains unexplored. In this work, we hypothesized that *Bari-Jheh* could also be affecting the expression of nearby genes by remodeling the local chromatin state. Thus, we tested whether *Bari-Jheh* is associated with H3K4me3, H3K9me3, and/or H3K27me3 histone marks, and whether stress affects this association. Finally, we also investigated whether stress affects the association of another transposon, which also belongs to the *Bari1* family, with the same histone marks.

## METHODS

### Prediction of PREs and TREs

We used the database JASPAR ^16^ with 95% threshold to predict the presence of PREs/TREs in the genomic region containing *Bari-Jeh* insertion and the three *Jheh* genes^17^. To identify PREs and TREs we used JASPAR matrix MA0255.1 and MA0205.1, respectively.

### Detection of piRNA reads

To search for piRNA homology sites in *Bari-Jheh*, we used the method described in Ullastres et al (2015)^18^. Briefly, reads were obtained from available piRNA libraries ^19,20^. We mapped the reads using BWA-MEM package version 0.7.5 a-r405 (Li 2013) to the 5.2 kb sequence including the *Bari-Jheh* element (chromosome 2R: 18,856,800-18,861,999). Then, we indexed and filtered sense/antisense reads by using samtools and bamtools ^21^. The total density of the reads was obtained using R (Rstudio v0.98.507).

### Detection of HP1a binding sites

To analyze the binding sited for HP1a in the *Bari-Jheh* region, we used HP1a modENCODE ChIP-Seq data ^22^, and we followed the methodology described above for mapping the reads to the *Bari-Jheh* region.

#### Fly stocks

We used the pair of outbred populations described in Guio et al. (2014)^12^ (DGRP #1), and a new pair of outbred populations created for this work (DGRP #2, Supplementary Table S1). Briefly, 10 virgin females and 10 males of seven strains homozygous for the presence of *Bari-Jheh* were placed in a large embryo collection chamber. The progeny was randomly mated during 10 generations before performing experiments. The same procedure was repeated for flies homozygous for the absence of *Bari-Jheh*. All the strains used to construct the outbred populations came from the Drosophila Genetic Reference Panel project ^23,24^. Flies were kept in large embryo collection chambers with regular fly food (yeast, glucose and wheat flour). Briefly, ∼200 mated females laid eggs during 24 hours. After that, the plates with eggs were placed in a new collection chamber and stored at 21-24°C until adult emergence. We selected adults emerged in a 24 hour interval and we placed them in a new chamber. After 48 hours, we split groups of 50 females with CO_2_ pads and store flies in tubes with fly food at 21-24°C until experiments were performed.

#### Oxidative stress exposure

To induce oxidative stress, we added paraquat to the fly food up to a final concentration of 10mM. For nonstress conditions, we used regular fly food. We transferred the flies to new tubes with or without paraquat and exposed them during 12 hours at 21-24°C before dissection. We did three to six replicas, of 50 females each, for each condition and genotype.

### Chromatin Immunoprecipitation assays

ChIP assays were performed with flies from two different genetic backgrounds. Three to six biological replicas of 50 flies each per background were analyzed. We performed ChIP assays in guts because the gut is the first barrier against oxidative stress ^25^. Guts of 5-day-old females were dissected in 1x PBS with protease inhibitor cocktail. After dissection, we homogenized the samples in Buffer A1 with a dounce tissue grinder (30 times). We crosslinked the guts with 1.8% formaldehyde during 10 minutes at room temperature. We stopped the crosslink adding glycine to a final concentration of 125mM. We incubated the samples 3 minutes and kept the samples on ice. We wash the samples 3 times with Buffer A1 and then we add 0.2ml of lysis buffer and incubate 3 hours at 4°C. After lysis, we sonicated the samples using Biorruptor^®^ pico sonication device from Diagenode: 15 cycles of 30 seconds ON, 30 seconds OFF. We used the Magna ChIP G chromatin immunoprecipation Kit (Millipore). Chromatin was immunoprecipitated with antibodies against H3K4me3 (Catalog # ab8580), H3K9me3 (#ab8898), and H3K27me3 (#ab6002). All the antibodies were ChIP grade and antibody quality was tested before performing the experiments (supplementary Figure 1). We separated 20µl for input and store it at −20°C. The remaining 180µl were divided in three aliquots and we added 1µl of each antibody plus 20µl of magnetic beads and dilution buffer up to 530 µl. We incubated the samples overnight at 4°C in an agitation wheel. After incubation we washed the beads with Low salt buffer, High salt buffer, LiCl complex buffer, and TE buffer. We separated the chromatin from the beads using 0.5 ml of elution buffer, including input samples. We added 1 µl of Proteinase K to each sample and incubated the samples at 65°C overnight in a shaker at 300 rpm. After incubation, we purified the samples using the columns provided by the kit. We stored the samples at −20°C until the qRT-PCR analyses were performed.

### qRT-PCR analysis

We quantify the IP enrichment by qRT-PCR normalizing the data using the “input” of each IP as the reference value, using the ΔCt method. Primers used for this study are described in Table S2. We first confirmed the quality of the antibodies and the specificity of the immunoprecipitation. We analyzed the enrichment of each histone mark in a well-known genomic region enriched for the different histone marks studied: *RpL32* (also known as *rp49*), *18SrRNA* (also known as *18S*), and *Ubx* enriched for H3K4me3, H3K9me3, and H3K27me3 respectively. We also tested whether the stress affects the histone marks enrichment in these three genes. We note that in a previous version of this work, we did not test whether stress affected the enrichment of histone marks in a well-known genomic region enriched for these marks ^26^. When we did this analysis, we found that stress did affect the enrichment on these genes suggesting that there were technical problems in the immunoprecipitation. We thus discarded these previous results.

### Statistical analysis

We used R software for the analyses. Results were not normally distributed. Different data transformation failed to normalize the data. Thus, we used a non-parametric test Kruskal-Wallis. Since we performed several tests we corrected the p-value for each set of tests using Bonferroni correction. We also tested whether results obtained with the two genetic backgrounds analyzed were significantly different before pooling them.

## RESULTS

### *Bari-Jheh* could be affecting the local chromatin state

To test whether *Bari-Jheh* could be affecting the local chromatin state, we analyzed the sequence of the transposon and its flanking regions including the three *Jheh* genes (Figure 1). We first looked for Trithorax group Response Elements (TREs) that recruit H3K4 methyltransferases, and Polycomb group Response Elements (PREs) that recruit H3K27 methyltransferases (see Material and Methods^27,28^. While H3K4me3 is associated with active promoters, H3K27me3 is associated with silenced or repressed promoters and enhancers ^29^. We found no TREs in the sequence analyzed, but we found one PRE in the *Bari-Jheh* sequence, and one PRE in the coding region of *Jheh3*, where modENCODE reports a Polycomb mediated repressive chromatin state (Figure 1A and 1B)^30^.

**Figure 1.**
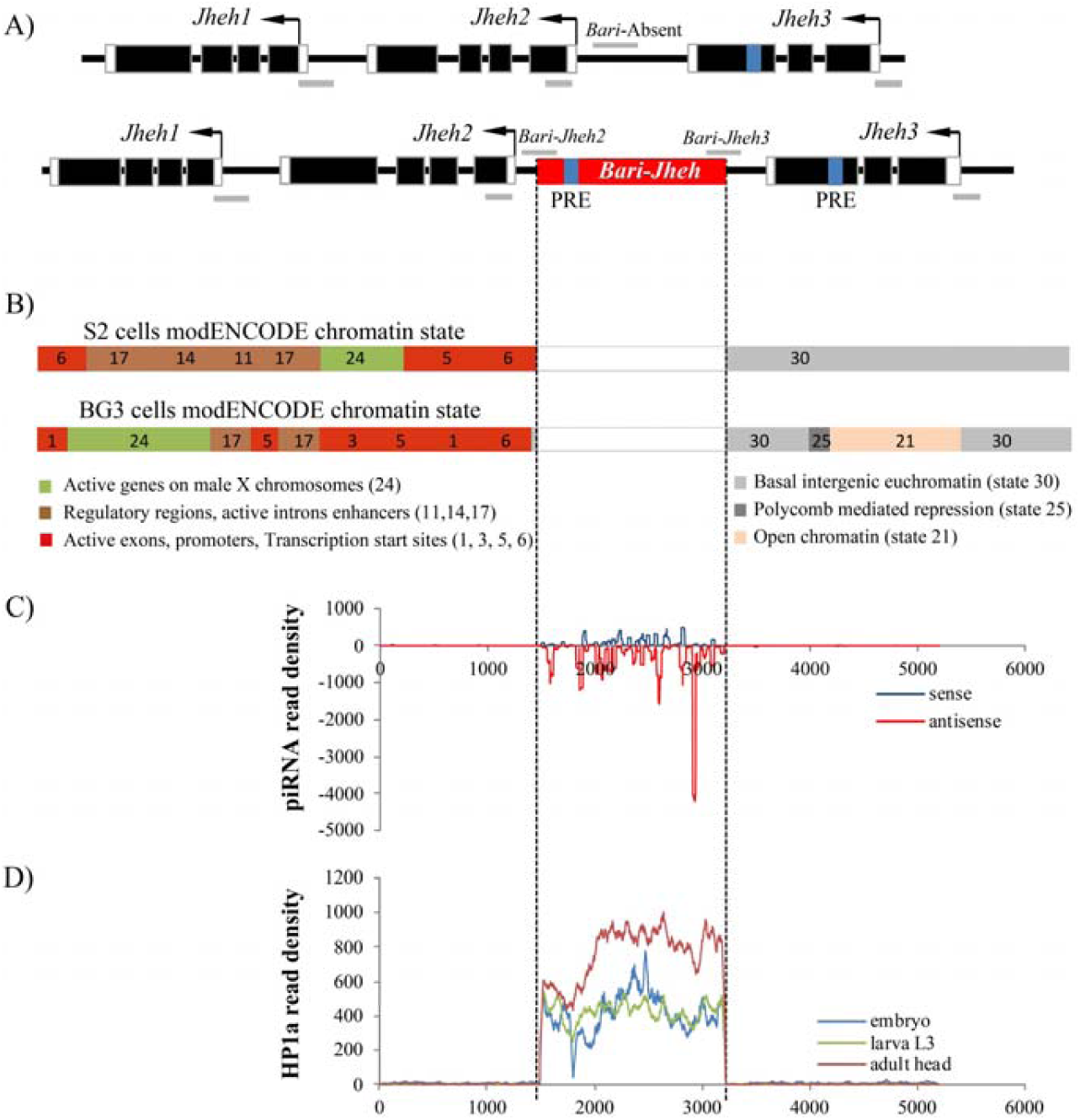
*Bari-Jheh* could be adding heterochromatin marks to the *Jheh* intergenic region. A) Schematic representation of *Jheh* genes in flies without *Bari-Jheh* and flies with *Bari-Jheh*. Black boxes represent exons, black arrows represent the direction of transcription, white boxes the 5’-UTR and 3’-UTR regions, the black line indicates intergenic or intronic regions and the red box represents *Bari-Jheh*. Grey lines represent the amplicons of the five regions analyzed using ChIP-qPCR experiments. The blue bars indicate the approximated position of the predicted PREs. B) modENCODE chromatin states in S2 cells and BG3 cells in the region analyzed. S2 cells and BG3 cells are derived from late male embryonic tissues and the central nervous system of male third instar larvae, respectively (Roy et al 2010). Colors and numbers represent different chromatin states. The vertical discontinuous lines indicate the location of *Bari-Jheh* insertion, that was not analyzed by modENCODE. C) Mapping of piRNA reads in the *Bari-Jheh* and flanking regions. Reads mapping in sense orientation are represented in blue, and reads mapping in antisense orientation in red. D) Mapping of HP1a reads in the *Bari-Jheh* and flanking regions. Reads from embryo stage are represented in blue, reads from larva L3 stage in green, and reads from adult head in red.

To further test whether *Bari-Jheh* affects the local heterochromatin state, we also investigated whether *Bari-Jheh* has piRNA binding sites and/or recruits HP1a (see Material and Methods). Sites with homology to piRNAs behave as cis-acting targets for heterochromatin assembly, which is associated with HP1a and H3K9me2/3 ^31^. H3K9me2/3 typically labels transcriptionally silent heterochromatic regions ^8^. We found that *Bari-Jheh* has sites with homology to piRNAs (Figure 1C). Accordingly, we also found that HP1a specifically binds to the *Bari-Jheh* sequence (Fig. 1D). Thus, *Bari-Jheh* could be introducing PREs that would be involved in the recruitment of H3K27me3 methyltransferase enzymes. Additionally, *Bari-Jheh* could also be inducing piRNA mediated heterochromatin assembly, and thus could be adding H3K9me2/3. These results provide suggestive but not conclusive evidence that *Bari-Jheh* could be introducing heterochromatin histone marks.

### *Bari-Jheh* is associated with an enrichment of H3K9me3 histone mark in nonstress conditions

To experimentally test whether *Bari-Jheh* affects histone marks enrichment, we performed Chromatin Immune Precipitation (ChIP)-qRT-PCR experiments in guts of adult flies with antibodies anti-H3K4me3, anti-H3K9me3, and anti-H3K27me3. We first tested the quality of the immunoprecipitation using genes previously reported to be enriched for the three histone marks analyzed (Figure S1, Table S3). We performed these analyses in flies with and without *Bari-Jheh* from two different genetic backgrounds, in nonstress and in stress conditions. As expected, we found significant enrichment of H3K9me3 for *18SrRNA* ^32^, significant enrichment of H3K4me3 for *RpL32* ^33^, and significant enrichment of H3K27me3 for *Ubx* ^34^ (Figure S1, Table S3). Note that *18SrRNA* is also enriched for H3K27me3. We did not find significant differences between strains with and without *Bari-Jheh*, or between nonstress and stress conditions (Figure S1, Table S3). Thus, we conclude that the immunoprecipitations were specific.

We compared the histone mark enrichment on both sides of *Bari-Jheh* insertion, *Bari-Jheh2* and *Bari-Jheh3* regions, with the corresponding region in flies without *Bari-Jheh*, *Bari*-Absent region (Figure 1). In nonstress conditions, we found significant differences in H3K9me3 in the *Bari-Jheh2* region compared with the *Bari*-Absent region in the two backgrounds analyzed (Figure 2). Thus, *Bari-Jheh* is associated with H3K9me3 enrichemnt in the *Jheh* intergenic region in nonstress conditions (Figure 2). Although we did not find an enrichment of H3K27me3 as expected from the presence of PRE elements in *Bari-Jheh,* our results are consistent with the presence of piRNA homology sites and HP1a in *Bari-Jheh* sequence (Figure 1).

**Figure 2.**
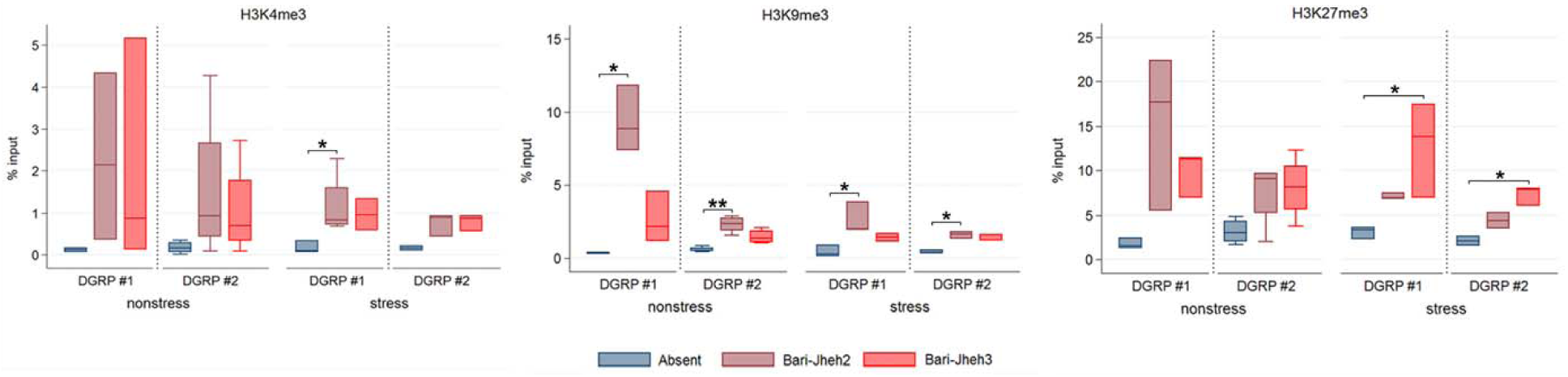
Histone mark enrichment in *Bari-absent, Bari-Jheh2* and *Bari-Jheh3* regions. Histone mark enrichment relative to the input of each strain for each background under nonstress and stress conditions. Each panel represents a different histone mark, H3K4me3, H3K9me3 and H3K27me3 in the *Bari-absent* (blue), *Bari-Jheh2* (light red) and *Bari-Jheh3* (red) analyzed regions. Significant differences between regions are mark with one (p-value <0.05) or two (p-value <0,01) asterisks.

### *Bari-Jheh* is also associated with an enrichment for H3K4me3 and H3K27me3 chromatin marks in oxidative stress conditions

To further test whether oxidative stress affects the chromatin marks added by *Bari-Jheh*, we performed ChIP-qPCR experiments in guts of adult flies exposed to paraquat. We found significant differences for H3K4me3 and H3K9me3 between flies with and without *Bari-Jheh*, in the *Bari-Jheh2* region in at least one of the backgrounds analyzed (Figure 2). We also found differences for H3K27me3 in the *Bari-Jheh3* region in the two backgrounds analyzed. Overall these results showed that in oxidative stress conditions, *Bari-Jheh* is associated with an enrichment of H3K4me3, H3K9me3, and H3K27me3 histone marks.

### *Bari-Jheh* did not affect histone marks enrichment on the nearby genes in nonstress conditions

Previous studies showed that spread of H3K9me3 histone mark to nearby DNA occurs at ≥ 50% of euchromatic TEs and can extend up to 20kb (average of 4.5 kb)^35^. Because we found that *Bari-Jheh* adds H3K9me3 in nonstress conditions, we tested whether there was also an enrichment of H3K9me3 in the promoter regions of the three genes nearby *Bari-Jheh* (Figure 1). In nonstress conditions, we found no significant enrichment for H3K9me3 in any of the three nearby genes *Jheh1*, *Jheh2* nor *Jheh3* (Figure 3). There were no differences either in the enrichment of H3K4me3 or H3K27me3 in the promoter regions of these three genes (Figure 3).

**Figure 3.**
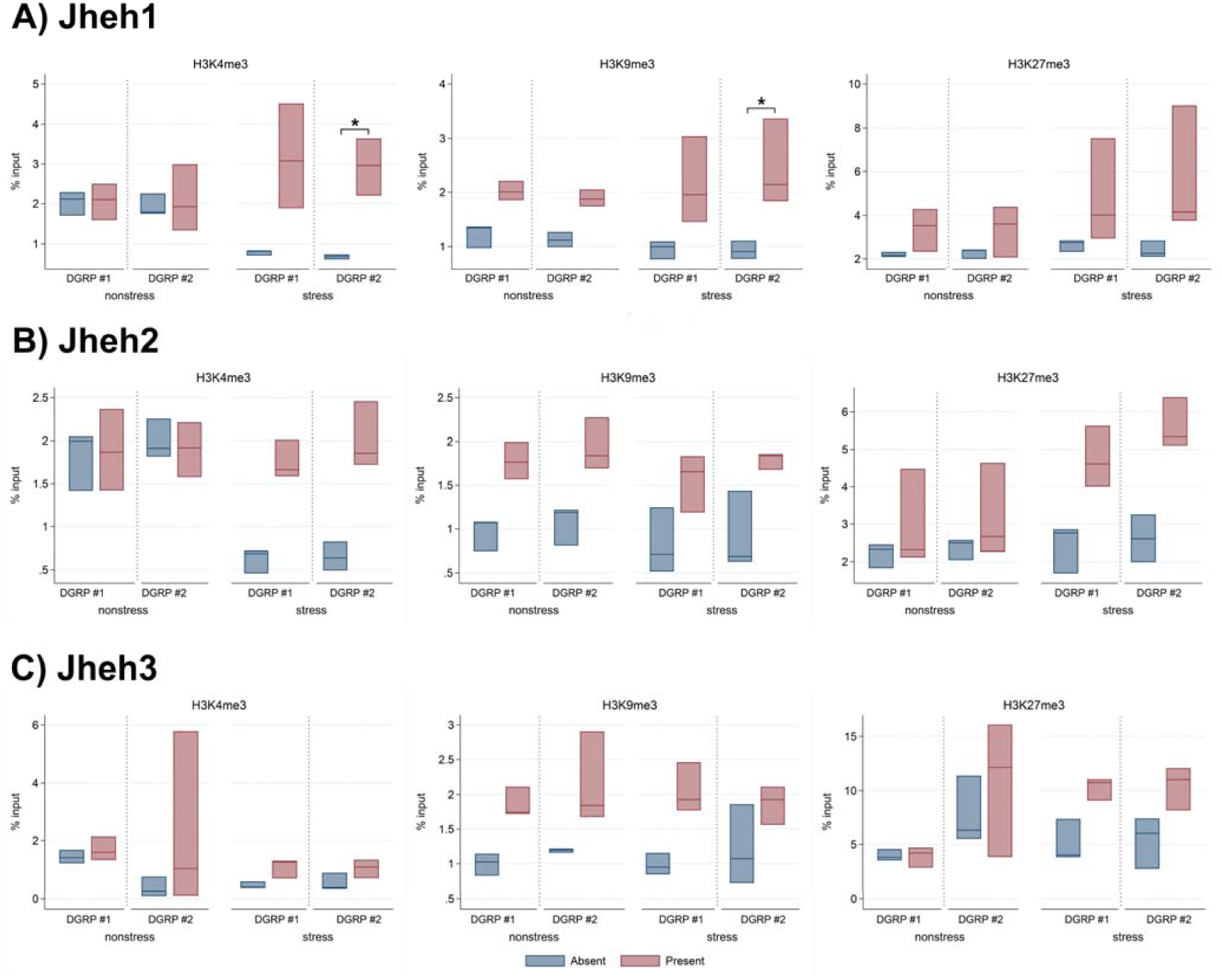
Histone mark enrichment in *Jheh1, Jheh2* and *Jheh3* gene up-stream regions. Enrichment of the histone marks relative to the input of each strain for each background under nonstress and stress conditions. Each panel represent a different histone mark, H3K4me3, H3K9me3 and H3K27me3 for A) *Jheh1*, B) *Jheh2* and C) *Jheh3* in the *Bari-absent* (blue) and *Bari-present* (light red) analyzed strains. Significant differences between regions are mark with an asterisk (p-value <0.05).

### *Bari-Jheh* is associated with H3K4me3 and H3K9me3 enrichment in *Jheh1* in oxidative stress conditions

We also tested whether we could detect a spread of H3K4me3, H3K9me3, or H3K27me3 to nearby DNA regions under oxidative stress conditions. We found that the promoter region of *Jheh1* showed enrichment for histone marks H3K4me3 and H3K9me3 in one of the backgrounds analyzed in strains with *Bari-Jheh* compared with strains without *Bari-Jheh* (Figure 3). On the other hand, no enrichment of H3K27me3 was detected in the regions nearby *Bari-Jheh* in any of the three genes analyzed (Figure 3). Thus, under oxidative stress conditions, we found enrichment of H3K4me3 and H3K9me3 in the promoter region of *Jheh1*.

### *Bari1-Cyp12a4* is enriched for H3K27me3 under oxidative stress conditions

We wanted to test whether other full-length insertions belonging to the *Bari1* family were associated with enrichment of histone marks. Besides *Bari-Jheh*, there are four full-length insertions located in the euchromatic region of the *D. melanogaster* genome. However, *FBti0019099* and *FBti0019419* were not present in the DGRP strains, and *FBti0019499* is flanked by other TE insertions, which precludes analyzing its presence/absence status. Only *FBti0019400* is fixed in the DGRP strains and thus we could study whether this copy was enriched for histone marks. *FBti0019400* is inserted in the 3’ UTR region of the cytochrome P450 gene *Cyp12a4*. The presence of *Bari1-Cyp12a4* insertion is associated with a shorter transcript and increased *Cyp12a4* expression ^36^. Because all the strains in our two pairs of outbred populations contain the *Bari1-Cyp12a4* insertion, we tested whether *Bari1-Cyp12a4* insertion showed different histone mark enrichment in nonstress *vs* stress conditions (Figure 4, Table 3). We did not find differences in the enrichment levels between the two backgrounds analyzed. Thus, we pooled the data from the two backgrounds. We found an enrichment of H3K27me3 when comparing nonstress *vs* oxidative stress conditions (Figure 4). Thus, we showed that *Bari1-Cyp12a4*, which also belongs to the *Bari1* TE family, showed an enrichment of H3K27me3 under oxidative stress conditions.

**Figure 4.**
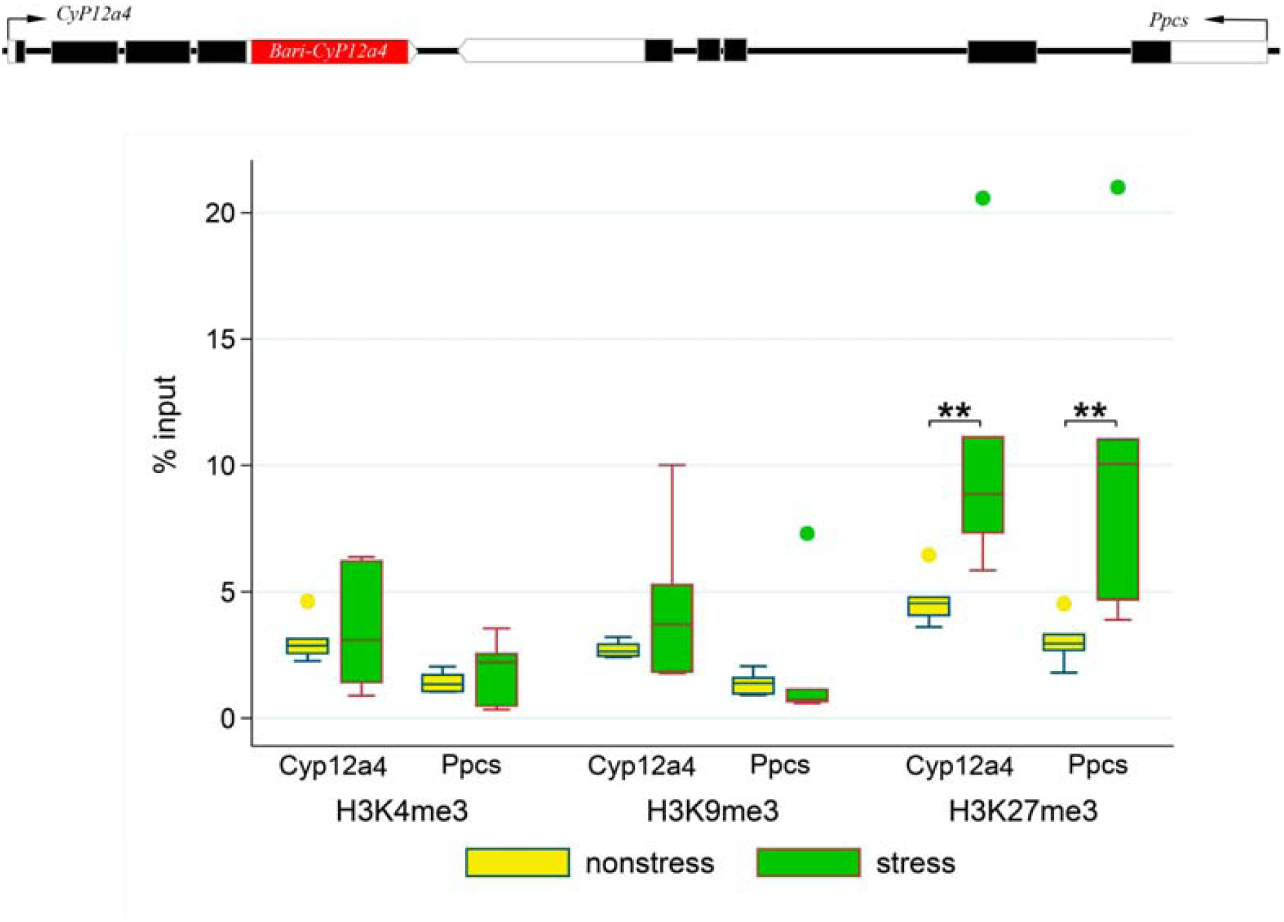
Histone mark enrichment in *Bari-Cyp12a4 and Bari-Ppcs* regions. Enrichment of the histone marks relative to the input of each *Bari* region for each histone mark under nonstress (yellow) and stress (green) conditions. Significant differences between regions are mark with an asterisk (p-value <0.05) or two (p-value <0,01).

**Table 1.** Statistical analyses of histone mark enrichment in the *Bari-Jheh* region. Significant values are highlighted in bold.

| Histone | Genetic Background | Treatment | Gene region | Comparison | K-W test p-value |
| --- | --- | --- | --- | --- | --- |
| H3K4me3 | DGRP #1 | Nonstress | *Bari-Jheh* | Abs – BJ2 | 0.1105 |
|  |  | Nonstress |  | Abs – BJ3 | 0.2696 |
|  |  | Stress |  | Abs – BJ2 | **0.0256** |
|  |  | Stress |  | Abs – BJ3 | 0.1516 |
|  | DGRP #2 | Nonstress | *Bari-Jheh* | Abs – BJ2 | 0.175 |
|  |  | Nonstress |  | Abs – BJ3 | 0.2547 |
|  |  | Stress |  | Abs – BJ2 | 0.0553 |
|  |  | Stress |  | Abs – BJ3 | 0.0789 |
| H3K9me3 | DGRP #1 | Nonstress | *Bari-Jheh* | Abs – BJ2 | **0.0109** |
|  |  | Nonstress |  | Abs – BJ3 | 0.2696 |
|  |  | Stress |  | Abs – BJ2 | **0.0109** |
|  |  | Stress |  | Abs – BJ3 | 0.2696 |
|  | DGRP #2 | Nonstress | *Bari-Jheh* | Abs – BJ2 | **0.0049** |
|  |  | Nonstress |  | Abs – BJ3 | 0.116 |
|  |  | Stress |  | Abs – BJ2 | **0.0169** |
|  |  | Stress |  | Abs – BJ3 | 0.2041 |
| H3K27me3 | DGRP #1 | Nonstress | *Bari-Jheh* | Abs – BJ2 | 0.038* |
|  |  | Nonstress |  | Abs – BJ3 | 0.115 |
|  |  | Stress |  | Abs – BJ2 | 0.2696 |
|  |  | Stress |  | Abs – BJ3 | **0.0109** |
|  | DGRP #2 | Nonstress | *Bari-Jheh* | Abs – BJ2 | 0.143 |
|  |  | Nonstress |  | Abs – BJ3 | 0.0937 |
|  |  | Stress |  | Abs – BJ2 | 0.2696 |
|  |  | Stress |  | Abs – BJ3 | **0.0109** |

**Table 2.** Statistical analyses of histone mark enrichment in the *Jheh* genes. Significant values are highlighted in bold.

| Histone | Genetic Background | Treatment | Gene region | Comparison | K-W test p-value |
| --- | --- | --- | --- | --- | --- |
| H3K4me3 | DGRP #1 | Nonstress | *Jheh1* | Abs – Pres | 1.00 |
|  |  | Stress |  | Abs – Pres | 0.0276* |
|  | DGRP #2 | Nonstress | *Jheh1* | Abs – Pres | 1.00 |
|  |  | Stress |  | Abs – Pres | **0.019** |
| H3K9me3 | DGRP #1 | Nonstress | *Jheh1* | Abs – Pres | 0.2101 |
|  |  | Stress |  | Abs – Pres | 0.0706 |
|  | DGRP #2 | Nonstress | *Jheh1* | Abs – Pres | 0.5227 |
|  |  | Stress |  | Abs – Pres | **0.0197** |
| H3K27me3 | DGRP #1 | Nonstress | *Jheh1* | Abs – Pres | 0.0944 |
|  |  | Stress |  | Abs – Pres | 0.4231 |
|  | DGRP #2 | Nonstress | *Jheh1* | Abs – Pres | 0.9245 |
|  |  | Stress |  | Abs – Pres | 0.1627 |
| H3K4me3 | DGRP #1 | Nonstress | *Jheh2* | Abs – Pres | 1.00 |
|  |  | Stress |  | Abs – Pres | 0.1627 |
|  | DGRP #2 | Nonstress | *Jheh2* | Abs – Pres | 1.00 |
|  |  | Stress |  | Abs – Pres | 0.1246 |
| H3K9me3 | DGRP #1 | Nonstress | *Jheh2* | Abs – Pres | 0.1246 |
|  |  | Stress |  | Abs – Pres | 0.0706 |
|  | DGRP #2 | Nonstress | *Jheh2* | Abs – Pres | 0.1246 |
|  |  | Stress |  | Abs – Pres | 0.1246 |
| H3K27me3 | DGRP #1 | Nonstress | *Jheh2* | Abs – Pres | 1.00 |
|  |  | Stress |  | Abs – Pres | 0.2101 |
|  | DGRP #2 | Nonstress | *Jheh2* | Abs – Pres | 1.00 |
|  |  | Stress |  | Abs – Pres | 0.1246 |
| H3K4me3 | DGRP #1 | Nonstress | *Jheh3* | Abs – Pres | 1.00 |
|  |  | Stress |  | Abs – Pres | 0.638 |
|  | DGRP #2 | Nonstress | *Jheh3* | Abs – Pres | 0.5277 |
|  |  | Stress |  | Abs – Pres | 0.7726 |
| H3K9me3 | DGRP #1 | Nonstress | *Jheh3* | Abs – Pres | 0.2101 |
|  |  | Stress |  | Abs – Pres | 0.0706 |
|  | DGRP #2 | Nonstress | *Jheh3* | Abs – Pres | 0.2683 |
|  |  | Stress |  | Abs – Pres | 0.2683 |
| H3K27me3 | DGRP #1 | Nonstress | *Jheh3* | Abs – Pres | 1.00 |
|  |  | Stress |  | Abs – Pres | 0.2683 |
|  | DGRP #2 | Nonstress | *Jheh3* | Abs – Pres | 1.00 |
|  |  | Stress |  | Abs – Pres | 0.3388 |

**Table 3.** Statistical analyses of histone mark enrichment in the *Bari-CyP* region. Significant values are highlighted in bold.

| Histone | Gene region | Comparison | K-W test p-value |
| --- | --- | --- | --- |
| H3K4me3 | *CyP12a4* | Stress - nonstress | 0.8728 |
|  | *Ppcs* | Stress - nonstress | 0.3367 |
| H3K9me3 | *CyP12a4* | Stress - nonstress | 0.6310 |
|  | *Ppcs* | Stress - nonstress | 0.1093 |
| H3K27me3 | *CyP12a4* | Stress - nonstress | **0.0065** |
|  | *Ppcs* | Stress - nonstress | **0.0065** |

## DISCUSSION

In nonstress conditions, *Bari-Jheh* adds H3K9me3 to the intergenic region between *Jheh2* and *Jheh3* genes (Figure 2 and Figure 5). H3K9me3 histone mark is associated with transcriptionally silent regions, and several TE insertions are enriched for this histone mark ^33,37,38^. The presence of this histone mark is consistent with changes in expression of *Bari-Jheh* nearby genes ^12-14^. Flies with *Bari-Jheh* showed lower levels of expression of *Jheh2* and *Jheh3* compared with flies without this insertion in nonstress conditions in DGRP#1 outbred population and in inbred strains, while no changes in expression were reported for *Jheh1* in the outbred population ^12-14^.

**Figure 5.**
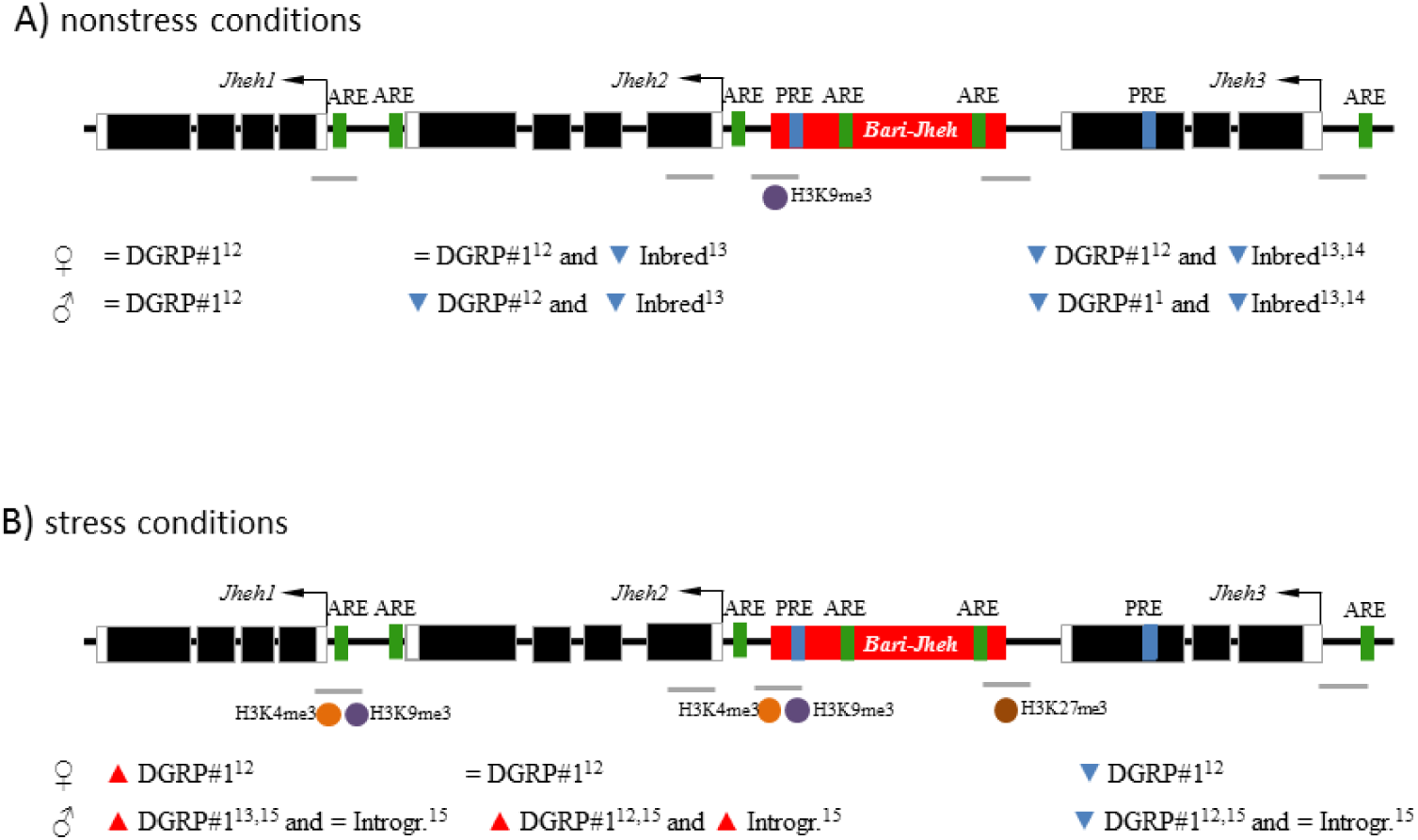
Summarize of findings. Representation of the histone mark enriched in each position for strains with *Bari-Jheh* compared with strains without *Bari-Jheh*. H3K4me3 is represented with an orange ball, H3K9me3 with a purple ball and H3K27me3 with a brown ball. Blue small triangles represent gene down-regulation, and red small triangles represent gene up-regulation. Upper scheme represents the enrichment under nonstress conditions and the lower scheme represents the enrichment under oxidative stress conditions.

We did not find evidence for the spreading of H3K9me3 found in the *Bari-Jheh* insertion to nearby genes as has been previously reported for ≥ 50% of euchromatic Tes 35. Consistently, no changes in expression in *Jheh1* have been found when comparing flies with and without *Bari-Jheh* insertion (Figure 5). Why some TEs are associated with the spreading of epigenetic marks while others are not is still an open question.

In oxidative stress conditions, the presence of *Bari-Jheh* is associated with enrichment of H3K4me3 and H3K9me3 both in the *Bari-Jheh2* intergenic region and in the promoter region of *Jheh1*. H3K4me3 and H3K9m3 have both been associated with active promoters ^29,39^. Indeed, both *Jheh2* and *Jheh1* genes have repeatedly been found to be up-regulated in flies with *Bari-Jheh* insertion under oxidative stress conditions 12,15. We also found that the *Bari-Jheh3* intergenic region is enriched for H3k27me3 in flies carrying the *Bari-Jheh* insertion. This result is consistent with the presence of a PRE in *Bari-Jheh* and in *Jheh3* gene. Consistently, we have previously found that *Jheh3* is downregulated in oxidative stress conditions in flies that carry *Bari-Jheh* insertion 12,15. In addition, we also found that *Bari1-Cyp12a4*, another TE insertion that belongs to the *Bari1* family, showed an enrichment of H3K27me3 under oxidative stress conditions Our results are thus consistent with previous findings in human cell culture that found an increase in the methylation marks in histone H3 in lysines K4, K27, and K9 in oxidative stress conditions ^40^.

Overall, our results suggest that besides adding cis-regulatory regions, *Bari-Jheh* also adds histone marks to the intergenic region between *Jheh2* and *Jheh3* genes, and it is associated with histone marks enrichment in the promoter of *Jheh1* gene. The presence of these histone marks was consistent with changes in expression previously reported for these three genes by analyzing flies with different genetic backgrounds differing in the presence/absence of *Bari-Jheh* insertion (Figure 5). How often the effect of TEs on gene expression is due to the presence of transcription factor binding sites in the TE sequence and/or to the enrichment of histone marks remains to be determined. Genome-wide analysis in which changes in gene expression are investigated together with binding of transcription factors and presence of histone marks in TE insertions in several genetic backgrounds are needed to solve this question.

## ACKNOWLEDGMENTS

We thank the *Equipe Eléments transposables, Evolution, Populations* for their support to L.G. We thank Elena Casacuberta for sharing antibodies and technical advice; Jon Frias for helping to create the new outbred populations, and Anna Ullastres for help with the piRNA and HP1a binding site analyses. We also thank the CNRS and the Molecular Biology facility DTAMB from the FRbioEnvis.

This work was supported by the Ministerio de Economia y Competitividad (BFU-2011-24397 and RYC-2010-07306 to J.G.); Secretaria d’Universitats i Recerca del Departament d’Economia i Coneixement de la Generalitat de Catalunya (2014SGR201 to J.G. and 2012FI-B-00676 to L.G.); European Commission (PEOPLE-2011-CIG-293860 and H2020-ERC-2014-CoG-647900 to J.G.) and Fondation pour la Recherche Médicale (DEP20131128536) and ANR Exhyb (ANR-14-CE19-0016) to C.V.

## AUTHORS CONTRIBUTION

L.G. designed the experiments, performed the experiments, analyzed the data and wrote the manuscript draft, C.V. and designed the experiments and analyzed the data and edited the manuscript, and J. G. designed the experiments, analyzed the data and wrote the manuscript.

## ADDITIONAL INFORMATION

“The authors declare no conflict of interest.”

Supplementary data in file “Supplementary data” with: Supplementary table S1: DGRP strains used for the study, Supplementary table S2: primers list used for the study, Supplementary table S3 p-values for positive control gene test. Supplementary figure S1: positive control genes.

## FIGURE LEGENDS

**Supplementary Figure 1: testing immunoprecipitations and antibodies.** Enrichment of the histone marks relative to the input of each strain for each background under nonstress and stress conditions. Each panel represents a different genomic region well known to enrich a different histone mark: A) *18SrRNA* enriched in H3K9me3 and H3K27me3, B) *RpL32* enriched for H3K4me3 and C) *Ubx* enriched for H3K27me3.

